# Sleeping Beauty transposon integrates into non-TA dinucleotides via an alternative mechanism

**DOI:** 10.1101/177584

**Authors:** Yabin Guo, Yin Zhang, Kaishun Hu

## Abstract

Sleeping Beauty transposon (SB) is an important genetic tool for generating mutations in vertebrates. It is well known that SB exclusively integrates into TA dinucleotides. However, this “TA law” has never been strictly tested in large number of insertion sites after next generation sequencing was widely utilized. In this study, we analyzed 600 million pairs of Illumina sequence reads and identified 28 thousand SB insertions in non-TA sites. We recovered some non-TA sites using PCR and confirmed that at least parts of the insertions at non-TA sites are real integrations. The consensus sequence of these non-TA sites showed an asymmetric pattern distinct from the symmetric pattern of the canonical TA sites. The right side of the consensus sequence is exactly the same as the sequence of SB transposon ends, indicating interaction between the transposon DNA and the target DNA. Based on these results we suggested that SB has an alternative integration mechanism besides the canonical one to integrate its DNA into non-TA sites.

**Highlights:** ∼ For the first time, we proved that Sleeping Beauty transposon can integrate into non-TA dinucleotides.

∼ For the first time, we provided evidence that transposon DNA can directly interact with target DNA.

∼ And for the first time, we found that a transposon can have two independent integration mechanisms.

## Introduction

The Sleeping Beauty transposon (SB) is a DNA transposon of Tc1/mariner family, which was constructed according to a consensus transposable element sequence from fish of Salmonid subfamily (1). SB is capable of transpose in mammalian system and is a popular genetic tool for generating genome-wide mutations in mammalian genomes (2–5). DNA transposons often have strong preference for their target sites. For example, PiggyBac strictly integrates into TTAA sites (6), while Hermes prefers T at the second position and A at the seventh position of its target site duplication (TSD) (7, 8). It is believed that SB (and all the other transposons of Tc1/mariner family) always integrates into TA dinucleotides based on the early results with limited integration events (9). In 2005, Yant *et al.* identified more than 1,300 SB integrations, which all targeted to TA dinucleotides (10). After next generation sequencing was utilized, even more SB integration sites were sequenced during the transposon mediated mutagenesis assays. The large number of integration events could have been an opportunity for further testing the “TA Law”. However, people already took for granted that TA dinucleotide is the only possible target for SB, so that sequence reads without a TA initial were considered artifacts and rejected in the sequence sorting and trimming before alignment (3, 4). Recently, an ever largest *ex vivo* SB mutagenesis screen was performed (5), in which more than 1,100 integration libraries were sequenced, 600 million pairs of sequence reads were obtained, and two million SB target positions were identified, providing a great convenience for mining rare integration events. In this study, we re-analyzed the sequence reads and found that SB did integrate into non-TA sites. And further analysis suggested that the SB insertions at non-TA sites are integrated via an alternative mechanism besides the canonical integrations mechanism.

## Results

### 28 thousand integrations at non-TA sites were identified by re-analyzing previous sequencing data

To search integrations in non-TA sites, we re-trimmed the sequence reads from previous study (5), accepting sequences with other initials as well as TA dinucleotide. After aligning to the mouse genome (mm10), 2,018,489 unique sites were identified, of which 28,794 insertions were at non-TA dinucleotides (Table S1, Fig. S1). Sequence reads from the same libraries that were aligned to the same coordinates and same strands were considered duplicates as previously described (11). Duplicates comprise sequence reads from independent insertions, cell duplications, as well as PCR amplifications. The insertions at non-TA sites were roughly 1.4% of the total insertions, and were termed as *general matches* in context (Table 1 and Fig. 1). To be more stringent, we assumed that those non-TA dinucleotides could be sequencing mistakes of TA dinucleotides. We replaced the first two nucleotides of the sequences with TA, and aligned them to the mouse genome again. All the sequences aligned successfully were removed from the *general matches* and resulted in a new assemble termed as *special matches*. The *special matches* are only slightly fewer than *general matches* (Table 1).

**Table 1:** Numbers of SB insertions identified at all of the 15 non-TA dinucleotides respectively

| Dinucleotides | General match | Special match |
| --- | --- | --- |
| CA | 5511 | 4916 |
| TG | 4853 | 3963 |
| TT | 2659 | 2366 |
| AA | 2600 | 2536 |
| GA | 2214 | 2168 |
| TC | 2046 | 1929 |
| AG | 1913 | 1907 |
| CT | 1478 | 1464 |
| GG | 1213 | 1211 |
| CC | 977 | 970 |
| AT | 894 | 890 |
| GT | 836 | 832 |
| AC | 767 | 750 |
| GC | 719 | 717 |
| CG | 114 | 112 |
| Total | 28794 | 26731 |

**Fig. 1.**
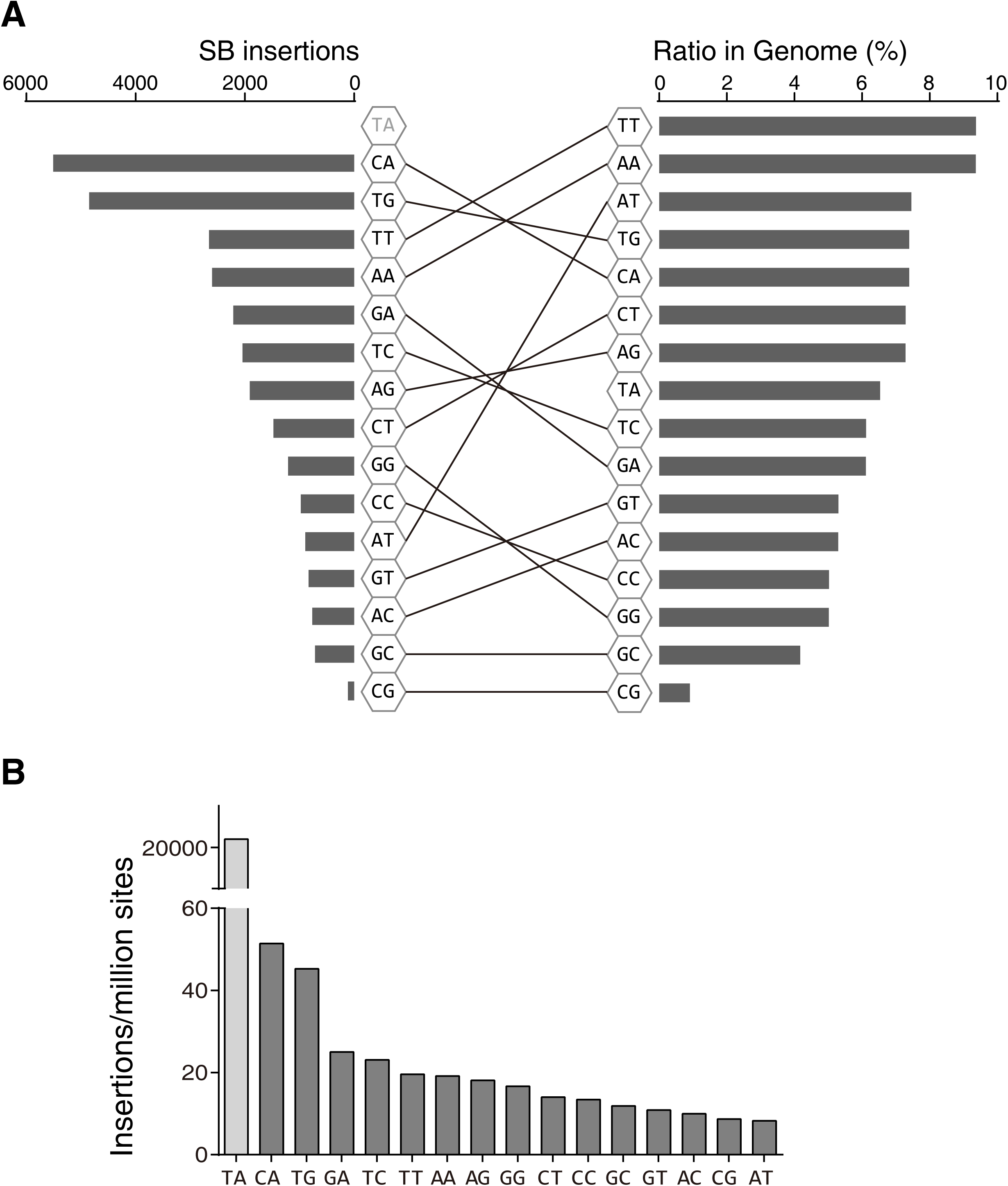
The SB insertions at non-TA dinucleotides and the ratio of dinucleotides in the mouse genome. Differential orders were shown between dinucleotides ranked by SB insertions and by their ratios in the mouse genome. B, SB insertions were normalized by the frequency of dinucleotides occurring in the mouse genome (only dinucleotides in non-repeat regions were calculated).

The integration frequencies at different dinucleotides are distinct. Considering the frequencies of the 16 dinucleotides are different in the mouse genome (Fig. 1 and Table S2), the distribution of insertions may be partially due to the distribution of dinucleotides in genome. For example, CG dinucleotides are very rare in mammalian genomes, and the insertions at CG dinucleotides are the fewest among all insertions. However, the order of nucleotides ranked by SB insertions is very different from that ranked by their occurrences in the mouse genome (Fig. 1A), indicating that some dinucleotides are more preferred by SB integration than others. The SB insertions were normalized by the dinucleotide frequencies in Fig 2B. TG/CA dinucleotides are apparently more preferred by SB integration than other non-TA dinucleotides, which may be because they are the most similar dinucleotides to TA (only one transition from TA).

**Fig. 2.**
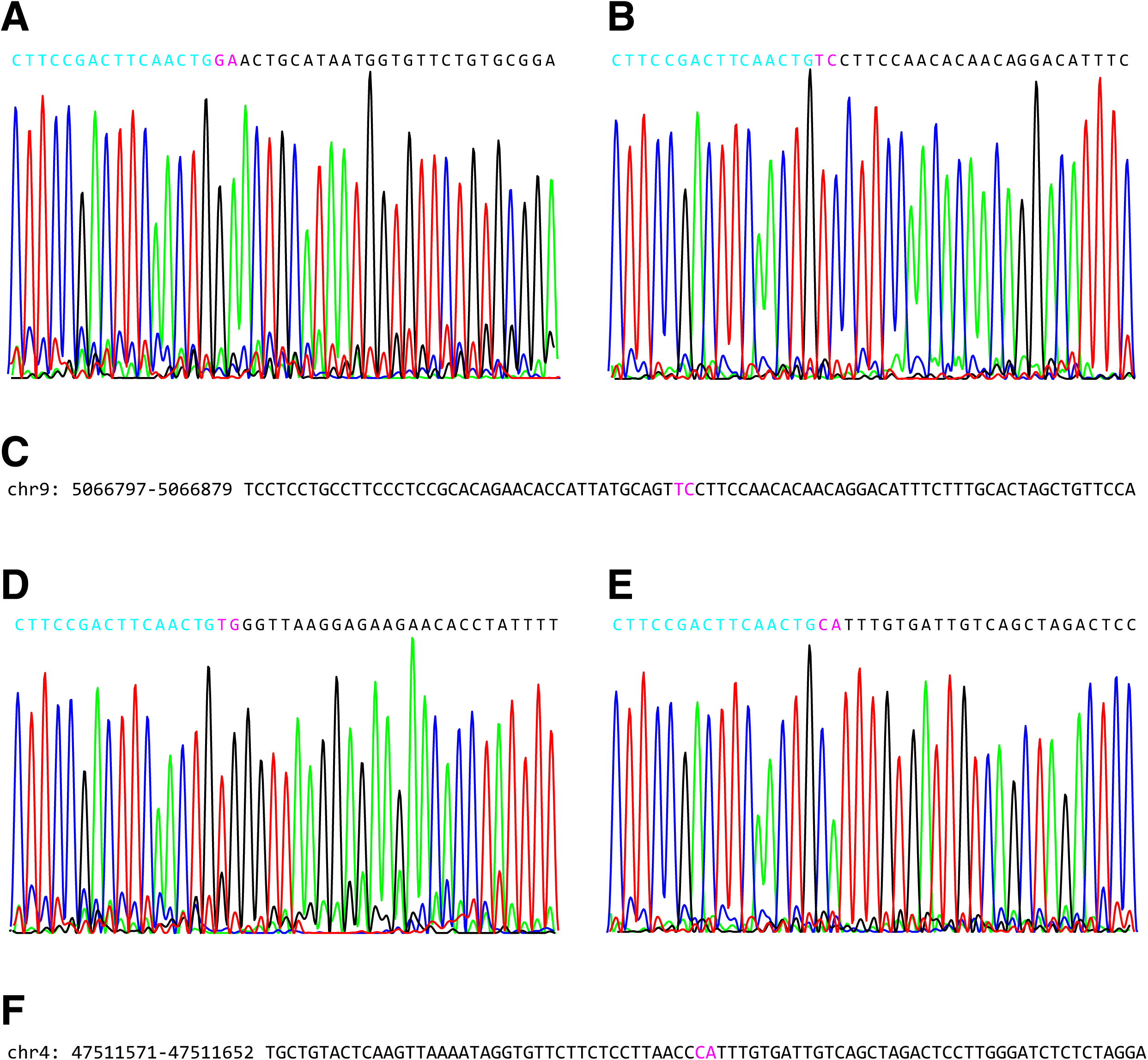
The recovery of SB integrations at non-TA sites. The SB integration site sequences were amplified from lib155.11 (A-C) and lib133.13 (D-E) using PCR. The cyan letters are the SB ends (A and D are right ends; B and E are left ends). The black letters are genomic sequences. The pink letters are the TSDs.

### Certain integrations at non-TA site were validated using PCR

To confirm whether the SB insertions at non-TA sites are real integrations or artifacts introduced by experiment design or data analysis, we picked nine insertion sites for PCR amplification. Since each library is a pool of cell clones with different integrations, only insertions with high duplicates were possible to be recovered (Table S1 and S3). For each insertion, two PCR reactions were performed. The primer pairs are [Primer 5, SB-left] and [SB-right, Primer 3] for insertions at plus strand, or [Primer 5, SB-right] and [SB-left, Primer 3] for insertions at minus strand (Table S3). Four out of nine insertions showed clear single bands in agarose gels for both primer pairs, which were then sequenced by Sanger sequencing. Fig. 2 showed two sites recovered from lib155.11 and lib133.13. Junctions between transposon and genomic sequences were found in both directions and perfect TSD was identified for each site (GA/TC for lib155.11, TG/CA for lib133.13; pink characters in Fig. 2A, B, D, E). This result indicated at least some of the non-TA sites found in sequence analysis are real integrations.

However, TSDs were not identified at certain sites (Fig. S2). The dinucleotides adjacent to the SB left ends are not TA, but the dinucleotides adjacent to the SB right end are still TA. This type of sites might be the result of *non-concerted* integrations, in which the TA dinucleotide is not duplicated. Because the LM-PCR for Illumina sequencing only detects the SB left end (5), the analysis can’t distinguish non-TA sites integrations from these *non-concerted* integrations. We examined the genomic sequences at all the non-TA sites and found that 1,922 sequences have TA right after the target dinucleotides, while 1,204 sequences have TA at the second position after target dinucleotides. Even if all these 3,126 sites were the results of non-concerted integrations, there are only a small part of the total insertions at non-TA sites. Therefore, non-concerted integrations couldn’t be a major contribution of the insertions at non-TA sites identified in our analysis.

### The target site sequences of non-TA sites have an asymmetric pattern

To show the sequence pattern of SB target sites, we extracted the SB target site sequences from the mouse genome according to their coordinates and strands. Sequence logos were then made based on these sequences (Fig. 3). Like many transposons/retrotransposons, the SB target site sequences show a perfect symmetric pattern (palindrome) (Fig. 3A). Recently, Kirk *et al.* stated that the palindromic consensus sequence at the target sites of some retroviruses is a result of integrations occurring “in approximately equal proportions on the plus strand and the minus strand of the host genome” (12). However, The symmetric sequence logos in some previous studies on transposon/retrotransposon (8, 13), as well as the present study, were all made of sequences with fixed orientation (*i.e.* reverse complement sequences were taken for integrations at minus strand). Therefore, palindromic sequences were favored at least by some transposon/retrotransposon integrations.

**Fig. 3.**
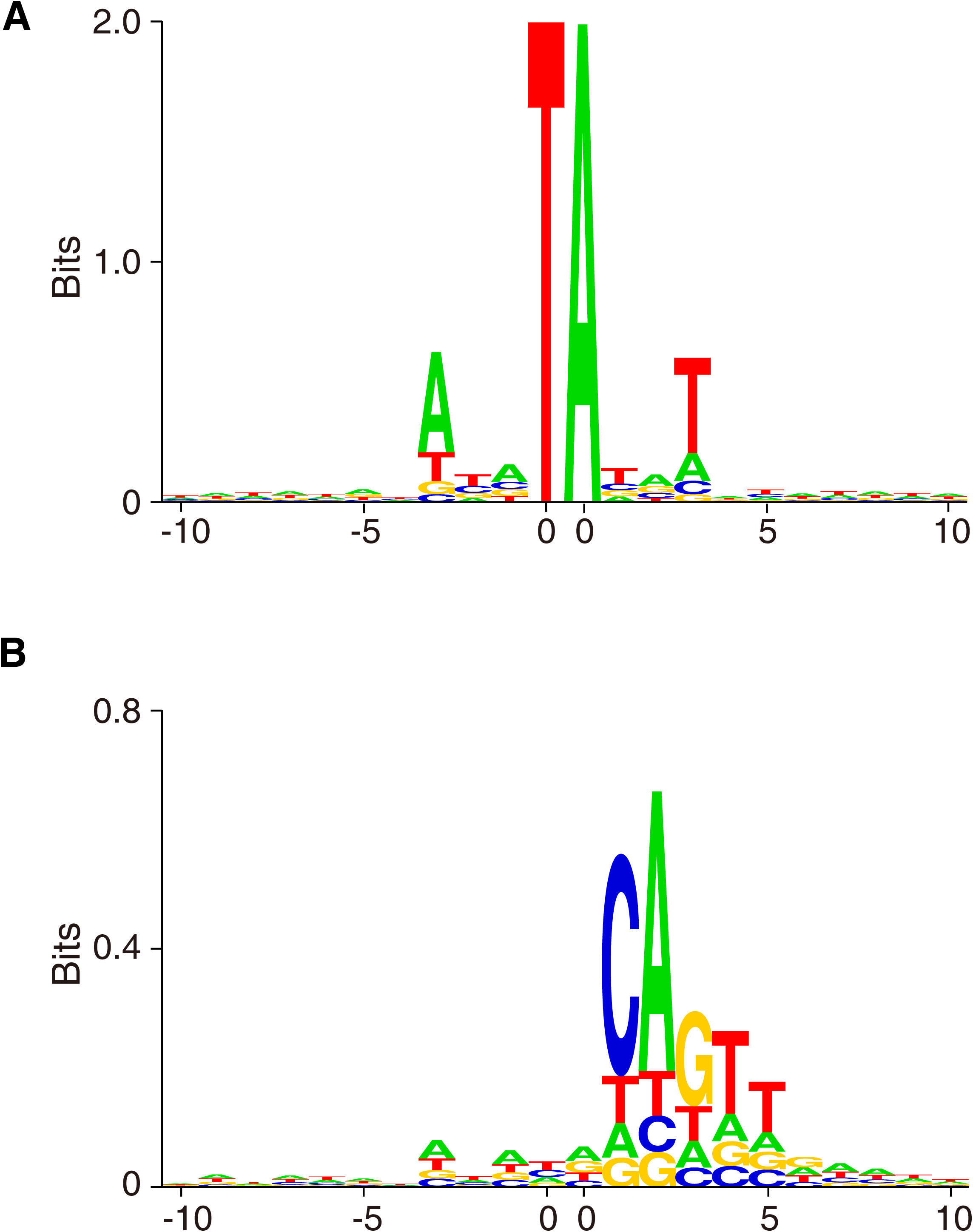
The sequence pattern in SB integration sites. The site sequences were aligned according to the SB target site and orientation. A, sequences of all target sites; B, sequences of non-TA target sites.

We then generated sequence logo with target site sequences only from non-TA sites. Strikingly, a distinct asymmetric pattern was revealed (Fig. 3B). The left side of the logo is basically the same as that of the canonical TA sites, whereas, the right side shows a conserved sequence, CAGTTGAA. Interestingly, this sequence is exactly the same as the SB transposon ends (Fig. S3). We also made sequence logos with sequences from different dinucleotides separately, and they all showed the similar pattern (Fig. S4-S6).

## Discussion

We re-trimmed the sequence reads, accepting all the other dinucleotides as well as TA dinucleotides, and identified 28 thousand SB targets at non-TA dinucleotides, which is around 1.4% of the total insertions. These insertions were not randomly distributed at non-TA dinucleotides. On the contrary, they showed preference between different dinucleotides, CA/TG being the most preferred ones (Fig. 1). To confirm if these integrations are real, we recovered some integration sites using PCR from the genomic DNA of some integration libraries. Sanger sequencing showed that two out of four sites have perfect TSDs, indicating they are real integrations (Fig. 2). Meantime, two *non-concerted* integrations were identified. We further employed some additional bioinformatics way to assess the ratio of *non-concerted* integrations. Finally, we speculate that the frequency of SB integrating into non-TA sites would roughly be around 1%.

To our knowledge, this is the first time that Sleeping Beauty transposon was proved to integrate into non-TA dinucleotides. There is still room for the well-established TA law to be improved. Although the ignorance of ∼1% integrations has little influence on the final analysis result, scientifically, we should know all the possibilities, no matter how small they are. Moreover, some recent studies had employed SB in gene therapy (14, 15), which has more risk concern than general molecular biology experiments. Rare integrations in non-TA sites may also need to be considered in gene therapies to minimize the unexpected insertions.

Unlike the general SB target sites, which have a palindromic consensus sequence, these non-TA sites show a distinct non-palindromic pattern (Fig. 3). The consensus sequence of the right side of the target site sequence (CAGTTGAA) is exactly the same sequence as the SB transposon end. We considered if the identical sequence pattern of the consensus sequence at target sites and the transposon end is an artifact: 1) the LM-PCR detected the junction between SB left end and genomic DNA, but the consensus sequence pattern is at the right side of target sites; 2) the target site sequences are extracted from the mouse genome, but not from sequence reads of Illumina sequencing; 3) there are no homologous sequences of SB in the mouse genome; even if the SB sequences were amplified in LM-PCR, they still couldn’t be aligned to the mouse genome (although there are CAGTTGAA in the mouse genome, it is not long enough for alignment); 4) the consensus sequence is a sequence of most frequent nucleotides, but is not necessarily a real sequence. Therefore, this sequence pattern couldn’t be an artifact.

Moreover, the insertions at non-TA sites are not homologous recombination, because: 1) the consensus sequence is not long and similar enough for homologous recombination; 2) the insertions have strong orientation bias (strand bias) (Fig. 3B); 3) the TSDs found in PCR (Fig. 2) indicated integration events.

We then focus on the 8-nucloetide box adjacent to the target dinucleotide at the right side of the target sites (assigned as R8 box for convenience), where the consensus sequence is located. Among the 28,794 non-TA sites, 12,732 (44%) R8 boxes are CAGnnnnn; 6,276 (22%) R8 boxes are CAGTTnnn; and 694 (2.4%) R8 boxes are CAGTTGAA. When the insertion numbers were normalized by the occurrence of the sequences in the mouse genome, we can see that the integration efficiency increased dramatically as the similarity between R8 box and the transposon end increased (Fig. S7). The frequency of SB integrated at a non-TA site adjacent to CAGTTGAA is >12,000/million sites, more than half of the integration frequency at canonical TA sites (Fig. 1B). Perhaps, that we saw many fewer integrations at non-TA sites is not because the integration efficiency at non-TA sites is low, but because there are many fewer CAGTTGAA sites than TA sites in the mouse genome.

Since the occurrence frequency for an 8-nucleotide sequence is roughly 4^-8^, the parity between these two sequences could not be a coincidence. Instead, it must be a result of complementarity between DNA strands. Therefore, we hypothesize that besides the canonical integration mechanism that relies on the interaction between transposase dimer/tetramer and TA dinucleotide, there is an *alternative integration mechanism* for SB transposon that relies on the interaction between the transposon DNA and the target DNA (Fig. 4). Due to the small contribution of the alternative integration to the entire integration collection, only when it is viewed separately from the canonical integration, can we notice its distinct property.

**Fig. 4.**
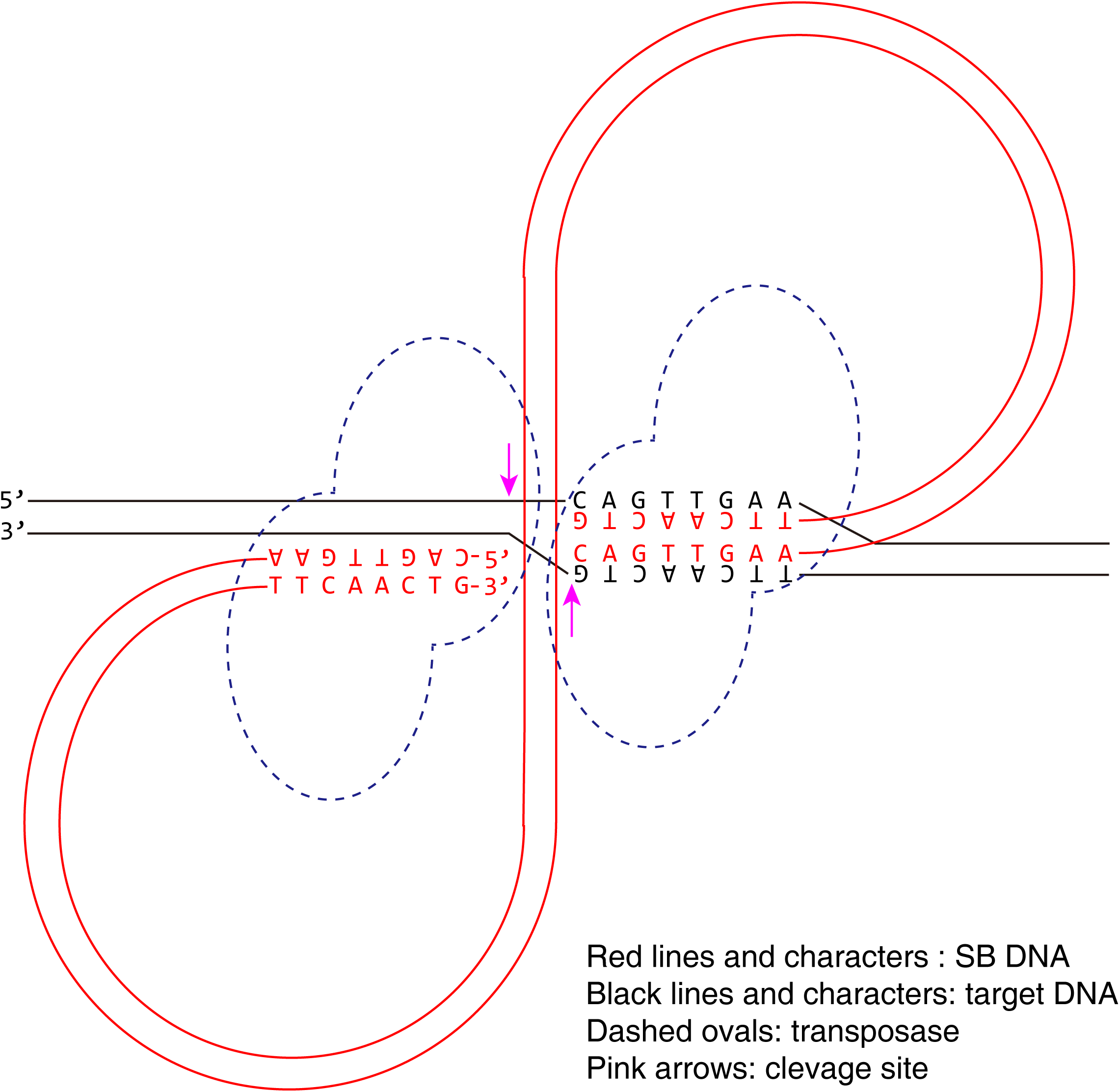
Model for the alternative integration mechanism of Sleeping Beauty transposon. The pre-integration complex (PIC) recognizes the target site via the interaction between the transposon DNA and the target DNA.

Why is there consensus sequence only at the right side of the target dinucleotides, but not the left side? We assume that because the IRDR-L and IRDR-R are somewhat different, after they bind to transposase, the conformations of two side of pre-integration complex (PIC) are different too, so that only the conformation of the right side is capable of interacting with target DNA strands.

Notably, that the alternative integration contributes for the insertion at non-TA sites doesn’t necessarily mean the alternative integration is the only source of the insertions at non-TA sites. The insertions at non-TA sites could be a pool of both canonical and alternative integrations, which is to be answered by future investigations.

In summary, we proved that SB transposon can integrate into non-TA sites. For the first time, we provided the evidence that transposon DNA can directly interact with target DNA to determine the integration sites. We also raised an idea that a transposon can have two independent integration mechanisms and provided a model for the alternative integration mechanism of SB transposon. In the future study, we will make point mutations to the SB transposon ends to see if the target site consensus sequence can be altered accordingly.

## Materials and methods

### Data source

The generation of the SB integration libraries in mouse BaF3 cells with SB100X transposase (16) and T2/Onc vector were described previously (5). The Illumina sequencing results were deposited in NCBI Short Read Archive, http://www.ncbi.nlm.nih.gov/sra. Accession no. SRX1491647.

### Bioinformatics analyses

Scripts for sequence trimming and dinucleotide frequency counting were written in Perl computer language. The trimmed sequences were aligned to the mouse genome using Bowtie 2 (17). The alignment output was filtered using a Perl script. The target site sequences were extracted from the mouse genome (mm10) using a Perl script. Reverse complement sequences were taken when the integration orientations are right-to-left (*i.e.* at minus strand). The target site sequence logos were generated using an application called DNAlogo (18), which had also been described in some previous studies (8, 13). The output PostScript (.ps) vector maps were converted to.pdf format in Adobe Illustrator.

### Recovering the SB target sites by PCR

Genomic DNA samples were extracted from the cell pools of SB integration libraries using DNeasy Blood & Tissue Kit (Qiagen), and were used as templates. Primers were designed according to the genomic sequences flanking the SB target sites and the SB left/right ends (Table S3). PCR reactions were performed using CloneAmp HiFi PCR Premix (Clontech). The PCR products were then sequenced by Sanger sequencing.

## Acknowledgement

We thank Dr. Kathryn O’Donnell of UT Southwestern Medical Center for her kind helps.

## Supplementary data

**Table S1.**

28,794 unique insertions at non-TA sites were identified. The chromosome, coordinate, strand, library, target site dinucleotide and total sequence reads were shown (see separated Excel file).

**Table S2.**
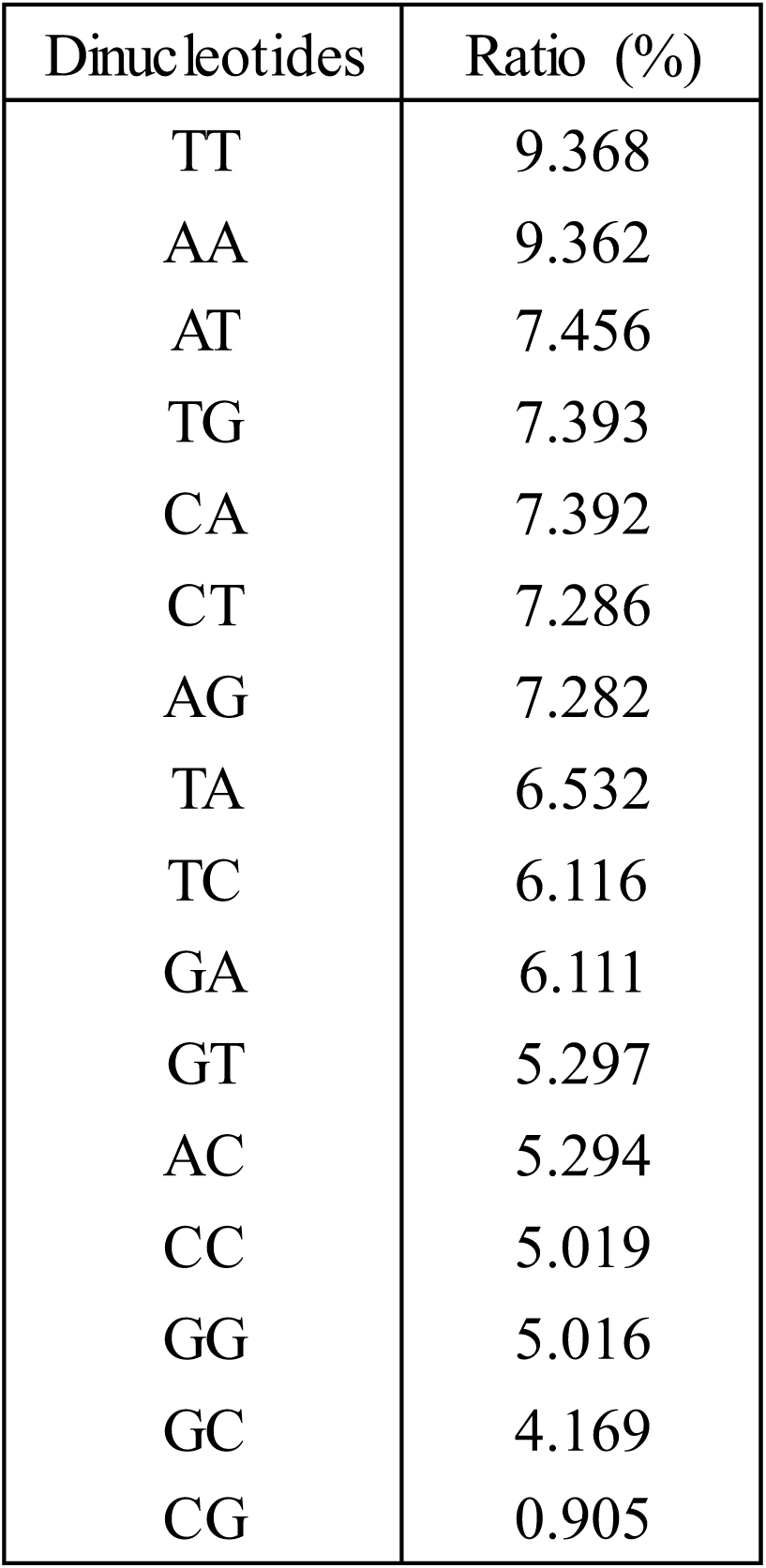
The ratios of all 16 dinucleotides occurring in the mouse genome (repeat regions were excluded, since the SB insertions in repeat regions could not be identified).

**Table S3.**

Target sites recovery using PCR. Non-TA sites with high duplicates were chosen. Primers were designed according to the genomic sequences flanking the target sites (see separated Excel file).

**Fig. S1.**
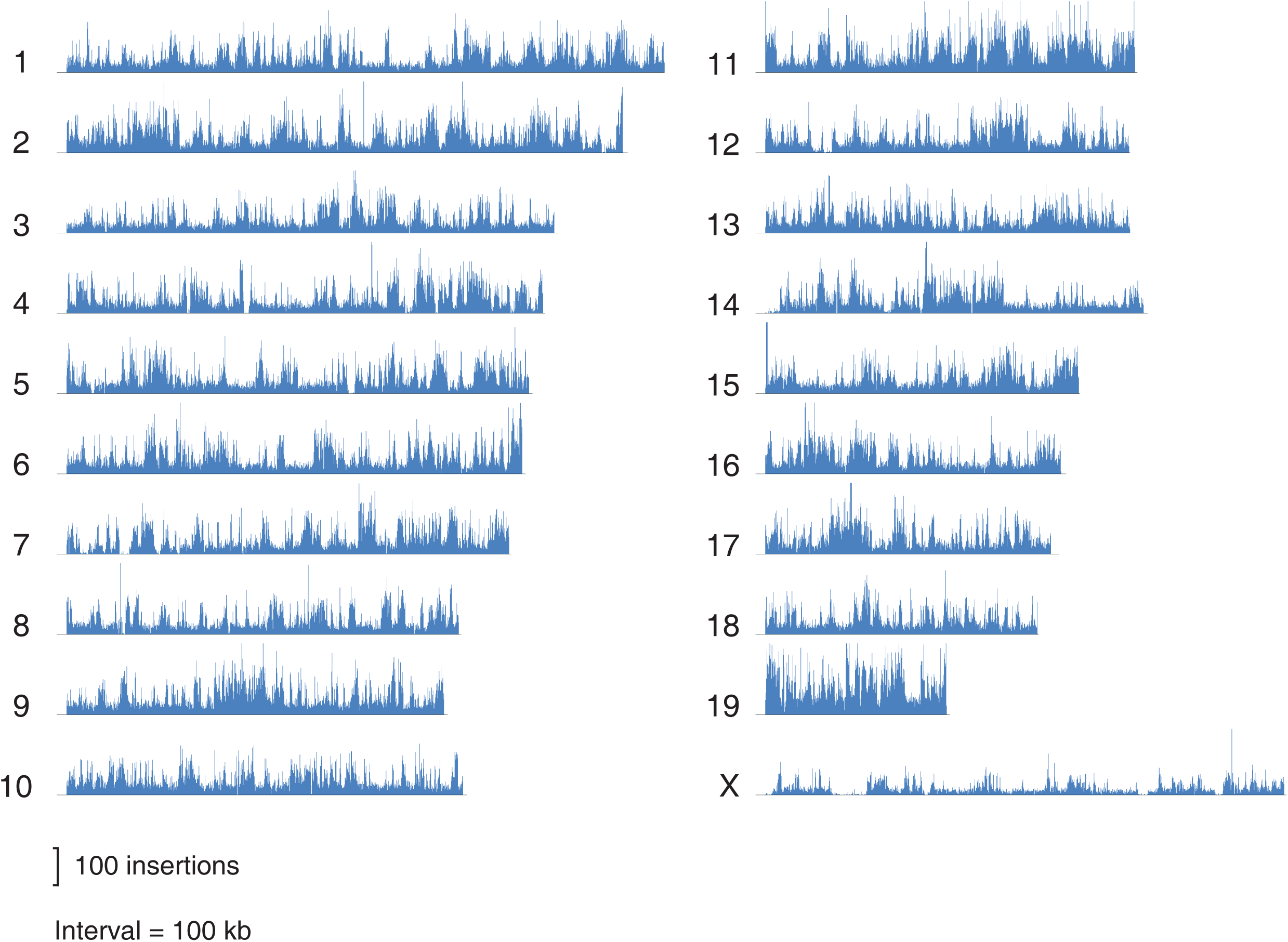
The distribution of 28,794 SB insertions on the mouse chromosomes. The insertions numbers per 100 kb interval were shown.

**Fig. S2.**
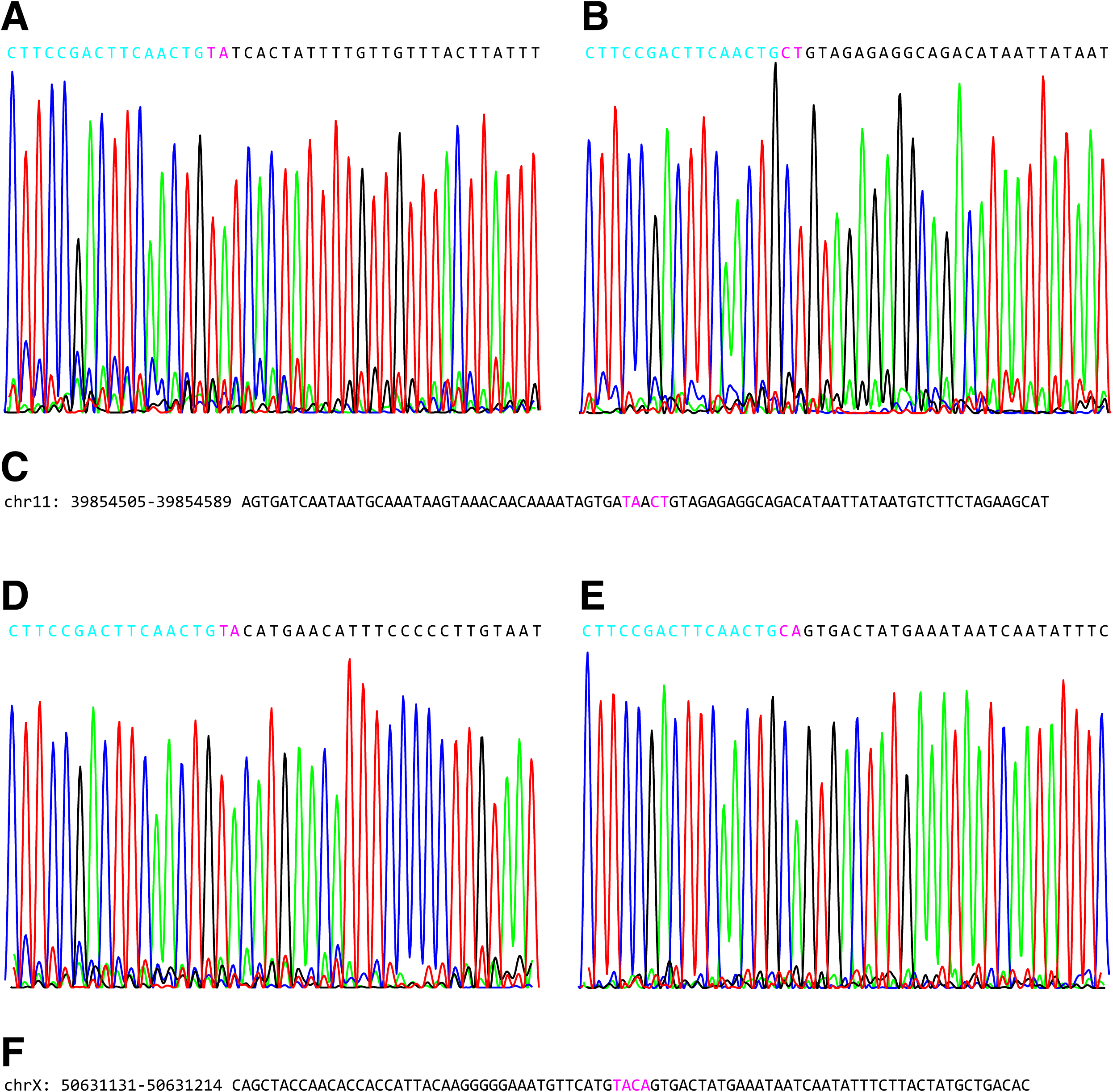
The identification of *non-concerted* SB integrations. The SB insertion site sequences were recovered from lib165.5 (A-C) and lib160.12 (D-E) using PCR. The cyan letters are the SB ends (A and D are right ends; B and E are left ends). The black letters are genomic sequences. The pink letters are the target site dinucleotides.

**Fig. S3.**
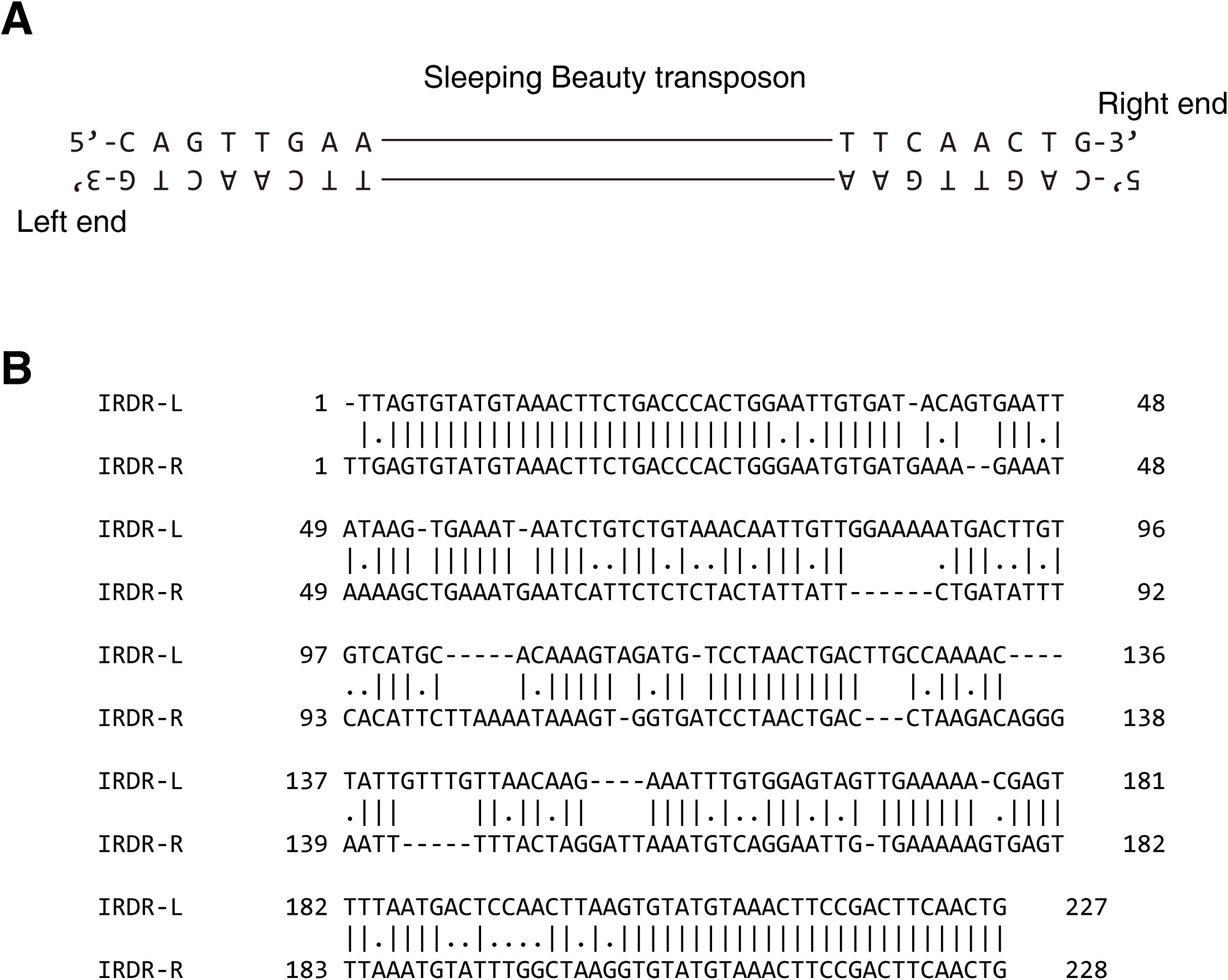
A, The sequences of left end and right end of the Sleeping Beauty transposon. B, The alignment of the SB IRDR-L and IRDR-R. IRDR, inverted repeat direct repeat.

**Fig. S4.**
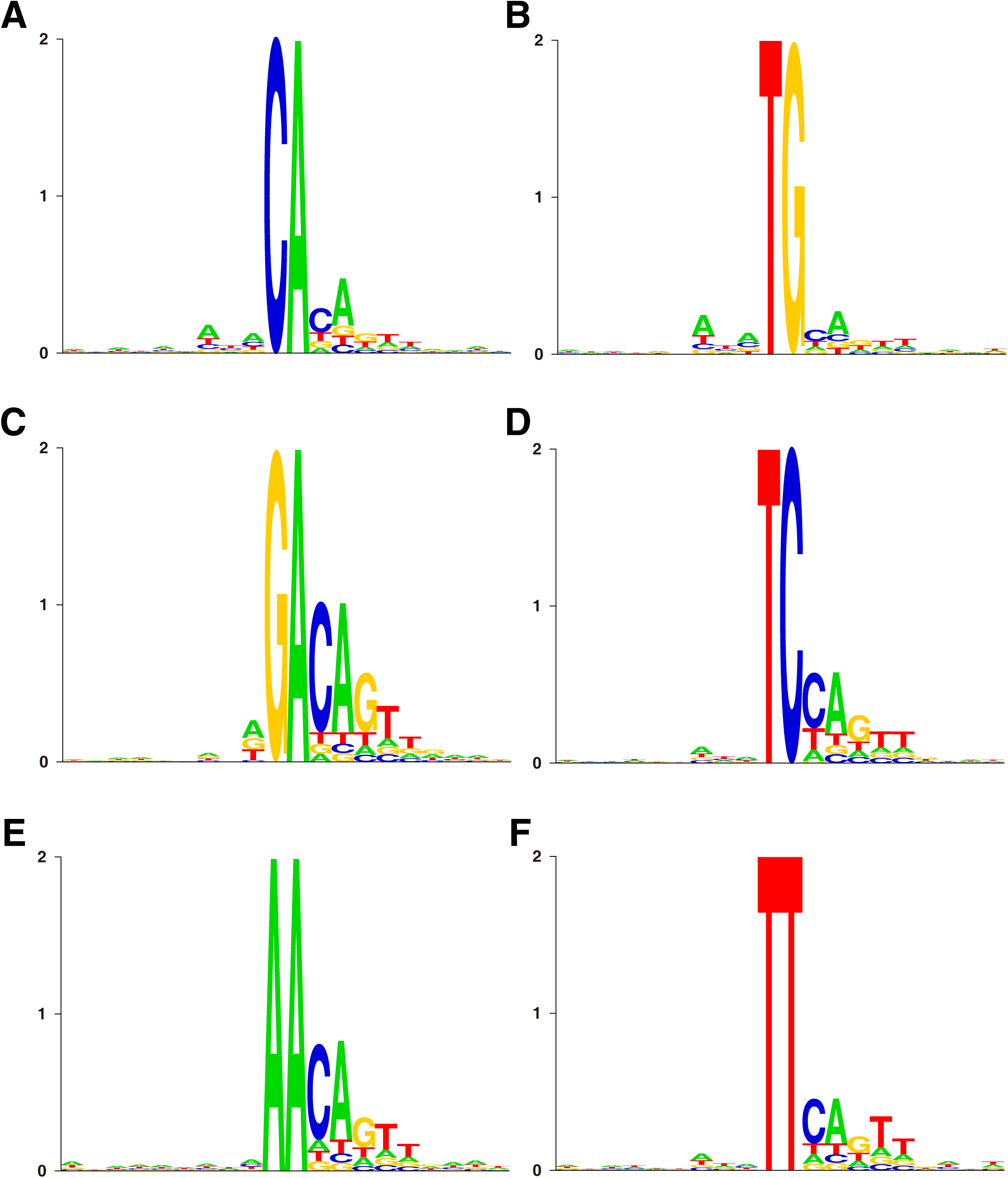
The sequence patterns at different non-TA target sites. A-F: CA, TG, GA, TC, AA, TT.

**Fig. S5.**
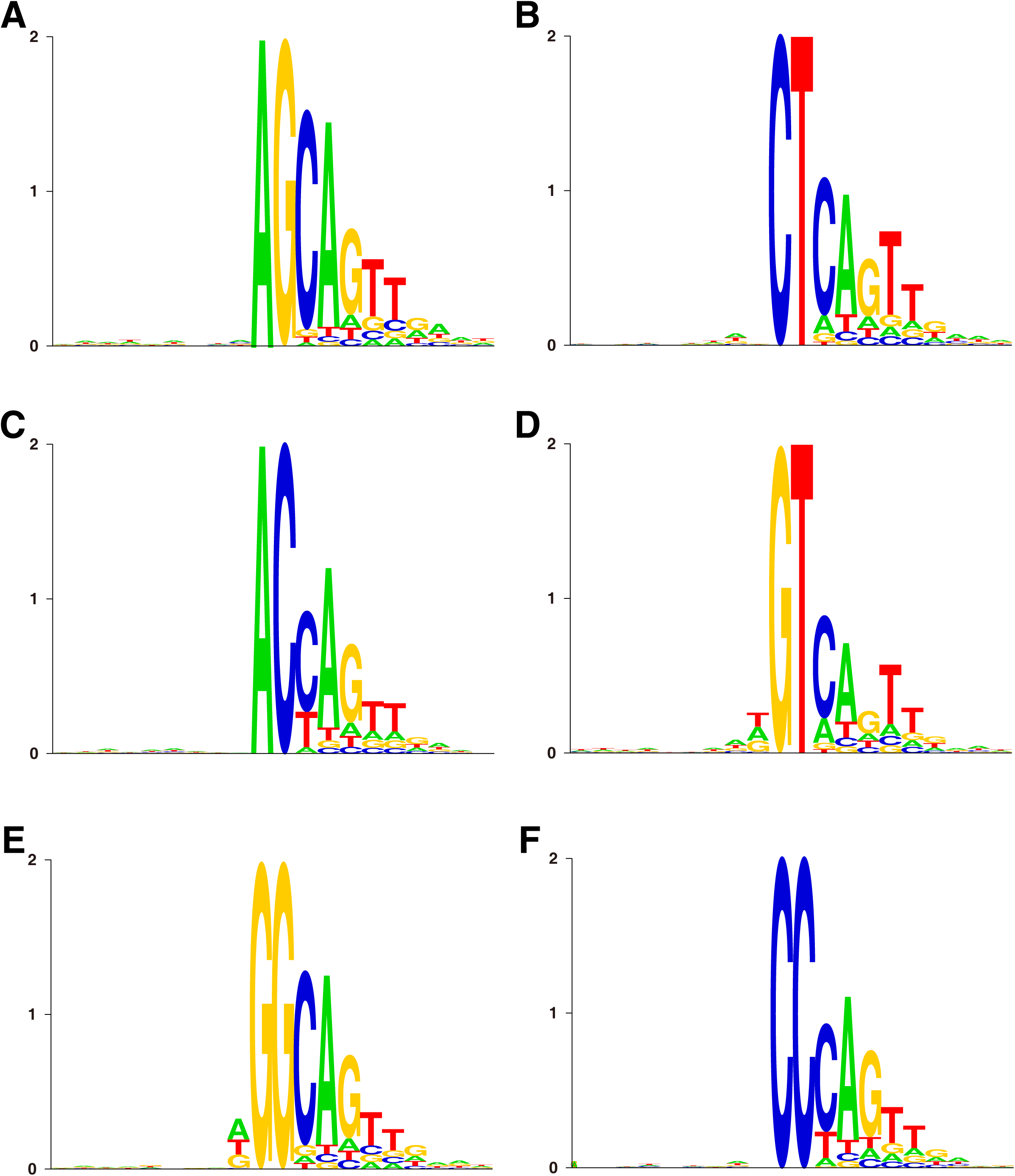
The sequence patterns at different non-TA target sites (continued). A-F: AG, CT, AC, GT, GG, CC.

**Fig. S6.**
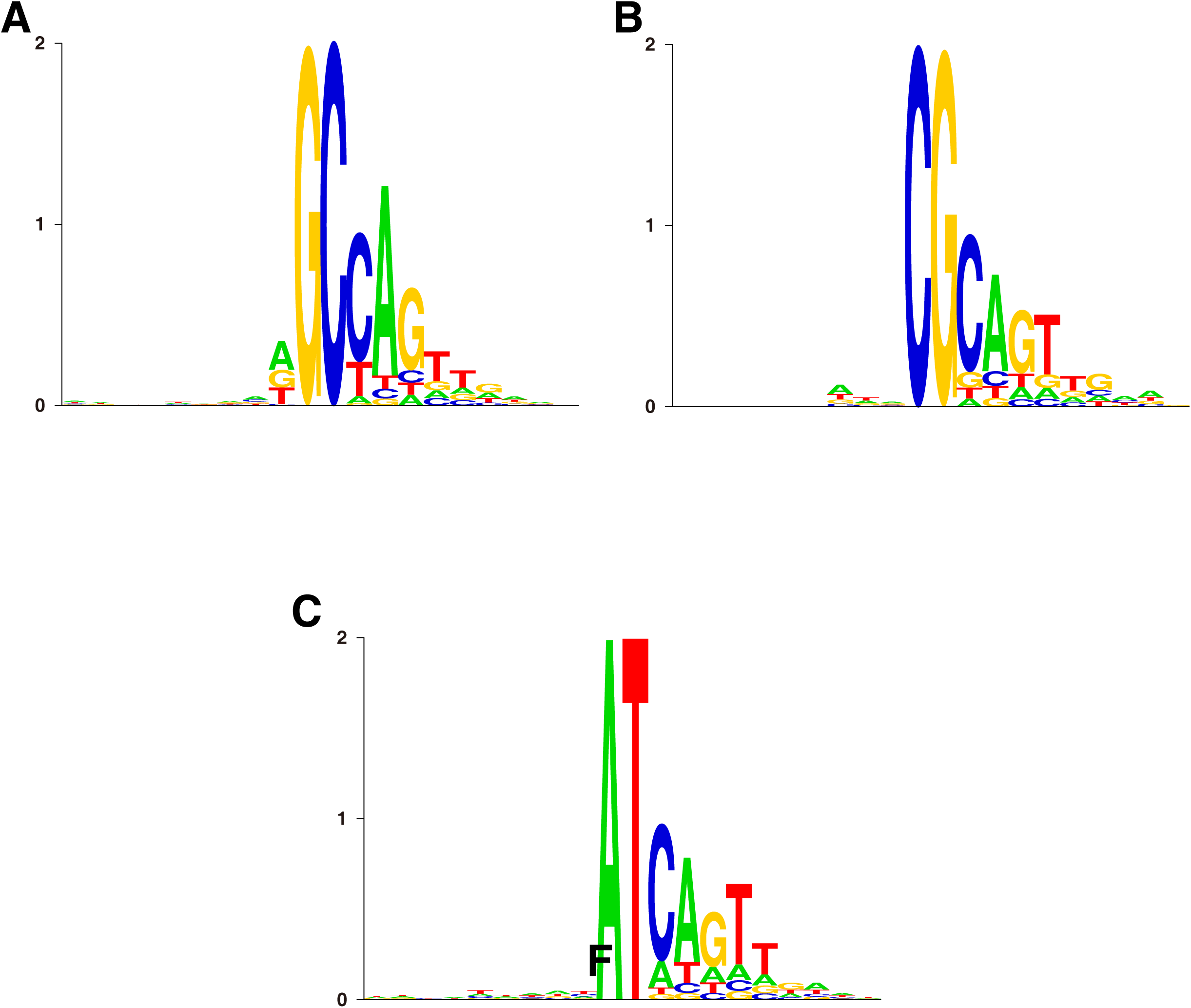
The sequence patterns at different non-TA target sites (continued). A-C: GC, CG, AT.

**Fig. S7.**
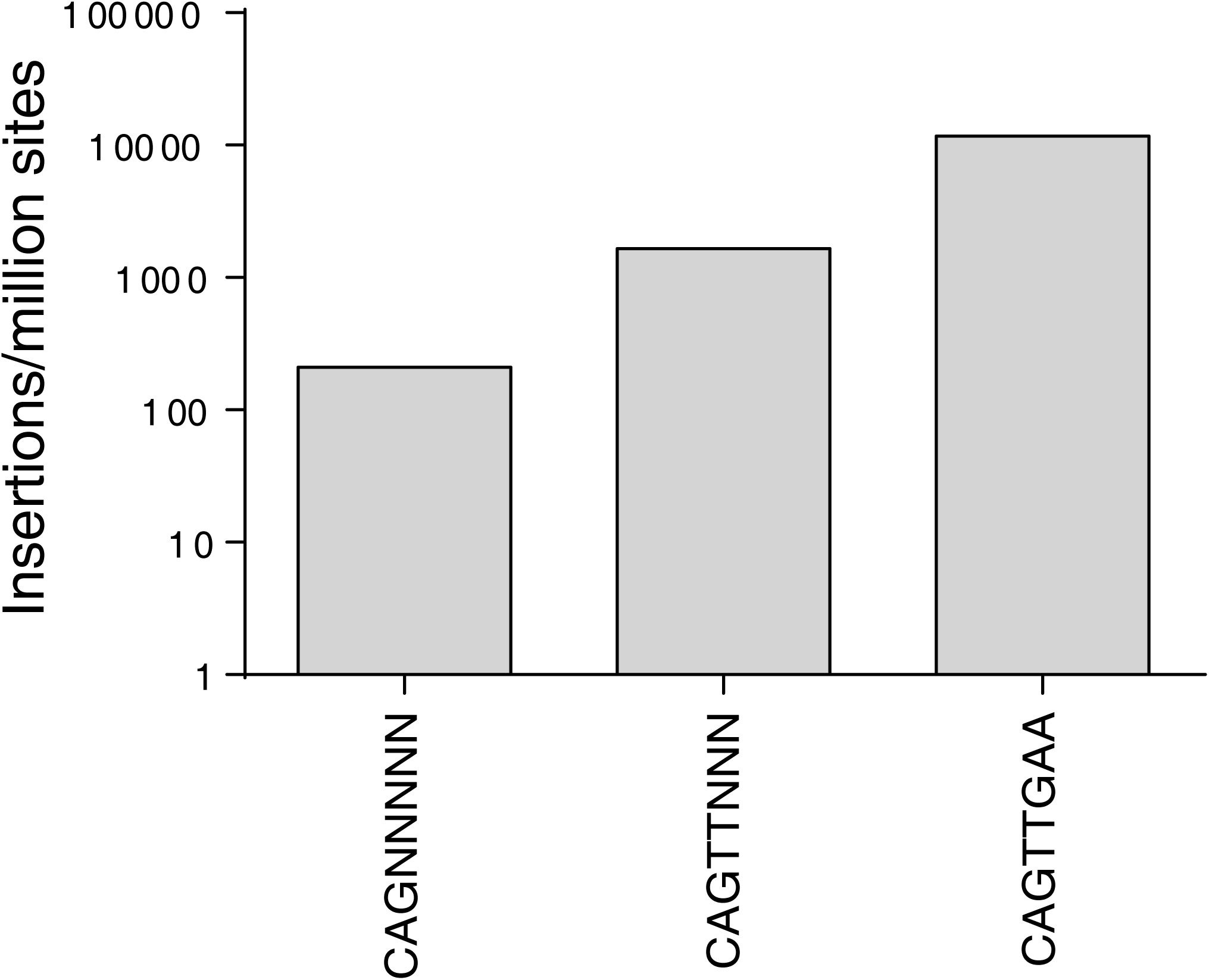
Number of SB insertions and R8 boxes. The number of SB insertions increased dramatically as the similarity between R8 box and the transposon end increased

